# pyHiM, a new open-source, multi-platform software package for spatial genomics based on multiplexed DNA-FISH imaging

**DOI:** 10.1101/2023.09.19.558412

**Authors:** Devos Xavier, Fiche Jean-Bernard, Bardou Marion, Messina Olivier, Houbron Christophe, Gurgo Julian, Schaeffer Marie, Götz Markus, Walter Thomas, Mueller Florian, Nollmann Marcelo

## Abstract

The three-dimensional (3D) nuclear organization of chromatin in eukaryotes plays a crucial role in gene regulation, DNA replication, and DNA damage repair. While genome-wide ensemble methods have enhanced our understanding of chromatin organization, they lack the ability to capture single-cell heterogeneity and preserve spatial information. To overcome these limitations, a new family of imaging-based methods has emerged, giving rise to the field of spatial genomics. In this study, we present pyHiM, an open-source and modular software toolbox specifically designed for the robust, automatic analysis of sequential spatial genomics data. pyHiM enables the reconstruction of chromatin traces in individual cells from raw, multicolor images, offering novel, robust and validated algorithms, extensive documentation, and tutorials. Its user-friendly graphical interface and command-line interface allow for easy installation and execution on various hardware platforms. The software employs a modular architecture, allowing independent execution of analysis steps and customization according to sample specificity and computing resources. pyHiM supports preprocessing, spot detection, mask detection, and trace generation, generating human-readable reports and intermediate results for data validation and further analysis. Moreover, it offers additional features for data formatting, result display, and post-processing. pyHiM’s scalability and parallelization capabilities enable the analysis of large, complex datasets in a reasonable time frame. Overall, pyHiM aims to facilitate the democratization and standardization of spatial genomics analysis, foster collaborative developments, and promote the growth of a user community to drive discoveries in the field of chromatin organization.

---

In eukaryotes, the three-dimensional (3D) nuclear organization of chromatin is tightly controlled and plays an active role in gene regulation, DNA replication and DNA damage repair. In the last decade, genome-wide ensemble methods, such as Hi-C and 3C ^1^, have revolutionized our understanding of genome structure at the kilobase-to-megabase scale by revealing the complex organization of chromatin into compartments, topologically-associating domains, and chromatin loops ^2,3^. However, these bulk approaches are unable to dissect single-cell heterogeneity or preserve spatial information in tissue ^4–6^.

Recently, a new family of imaging-based methods was developed to trace the 3D conformation of chromatin in single cells, giving rise to the field of spatial genomics ^7–11^ (Fig. 1a). These techniques perform sequential imaging of genomic loci with a precision of a few tens of nanometers, allowing for the 3D mapping of a given region of chromatin at kilobase resolution in thousands of individual cells ^8,9,12^. Our specific implementation, called Hi-M, couples detection of chromatin structure and transcriptional output ^8^ (Fig. 1a). Since their creation, spatial genomics methods based on sequential imaging were successfully used for the detection of short- and long-range chromatin interactions in multiple model systems, including mammalian cultured cells, fly embryos, and mouse tissues ^*7–10,12*^. Critically, imaging-based spatial genomics technologies complement transcriptomic surveys of single cells in their spatial context and thus have the potential to lead to important new discoveries in multiple fields, including 3D genomics, transcriptional regulation, DNA replication, or DNA repair.

**Figure 1.**
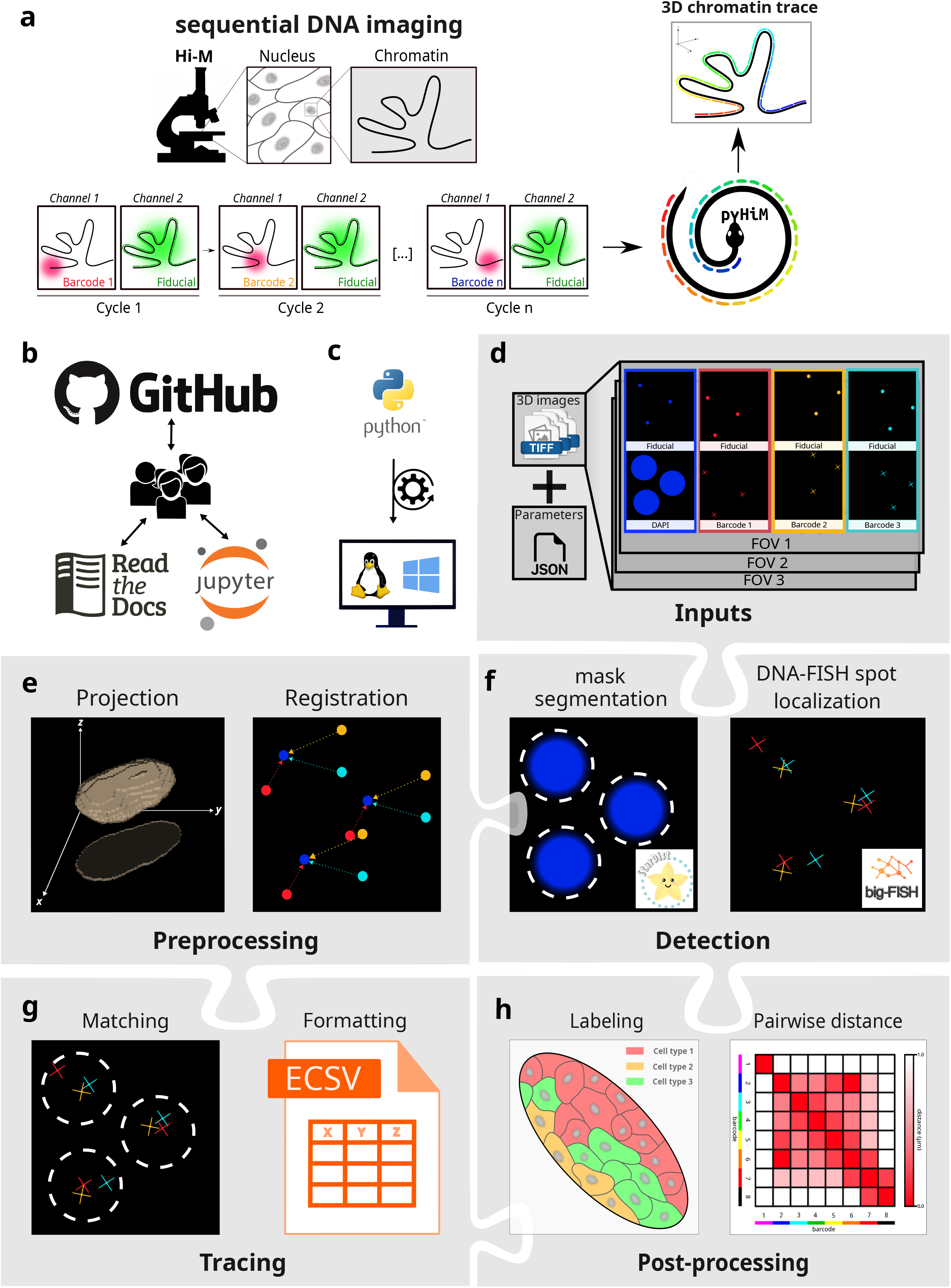
**a-** Schematic description of Hi-M microscopy: Chromatin is imaged through multiple acquisition cycles, each targeting a specific genomic locus using a set of unique DNA-FISH oligonucleotides targeted by a complementary, fluorescently-labeled oligonucleotide. A fiducial marker is simultaneously imaged to allow for registration and drift correction during post-processing. Using pyHiM, the 3D conformation of the target locus is reconstructed for each individual cell. **b-** pyHiM is an open source project hosted on GitHub. Extensive documentation and Jupyter notebooks are available for users and developers. **c-** pyHiM is developed in Python and runs indifferently on Linux and Windows. **d-** Input data: 3D images are organized by imaging channel (DAPI, fiducial, DNA-FISH spots, etc.) and FOV. A single json file combines all parameters needed to run the analysis pipeline. **e-** 3D images are pre-processed by calculating the maximum intensity projection and applying 2D registration based on the fiducial images. **f-** Masks for nuclei, oligopaint libraries, and DNA-FISH spots are computed using pre-trained deep learning models. Individual DNA-FISH spots are localized with sub-pixel accuracy using big-FISH. **g-** Individual traces are built by combining the localizations of all DNA-FISH spots detected within the same mask. Results are saved in ECSV format. **f-** Post-processing analyses are performed to obtain pairwise distance and proximity frequency matrices for each combination of DNA loci, and for different spatial regions of the sample containing different cell types.

In recent years, several efforts were made to promote a wider use of these new technologies by sharing experimental and image analysis protocols ^7,13–16^. However, democratization of spatial genomics will require the development of open-source and user-friendly software packages for reconstructing chromatin traces (i.e. unique sets of 3D coordinates describing a locus conformation in an individual cell) from raw, 3D, multicolor images ^17^. To this end, such software should: 1) provide access to validated cutting-edge techniques required for the analysis of spatial genomics data, 2) use a license-free programming language, 3) provide extensive documentation and tutorials to guide new users and allow development of new functionalities, 4) adopt a modular architecture to facilitate adaptation to future developments in spatial genomics, and 5) use novel analysis methods to ensure robust, automatic analysis of large data sets (several Tb per experiment) without user input in reasonable times.

To address these needs, we introduce pyHiM, an open-source, modular and scalable software toolbox specifically designed for sequential spatial genomics data analysis (Fig. 1a). pyHiM comes with extensive [user and developer documentation](https://pyhim.readthedocs.io/en/latest/index.html), as well as tutorials that illustrate typical analysis pipelines (Fig. 1b). It can be easily installed using standard package management tools ([conda](https://docs.conda.io/en/latest/) and [PyPi](https://pypi.org/), Fig. S1), and conveniently runs in both Windows and Linux (Fig. 1c). A single human-readable configuration file is used to centralize all analysis parameters and can be edited thanks to a user-friendly graphical user interface (GUI) (Fig. S1). In addition, a command-line interface enables execution on multiple hardware platforms, from laptop computers to high-performance computing (HPC) clusters. Functionality can be tuned according to local hardware specifications, acquisition conditions (*e*.*g*. number of channels, size of 3D image stacks), and sample properties.

The analysis pipeline of pyHiM is organized in modules, each performing a specific analysis task. The inputs of pyHiM are 3D image stacks in the universal TIFF format (Figs. 1a, 1d). Deconvolution of images before pyHiM execution is not mandatory for pyHiM analysis but, in our experience, improves the quality of the results and the statistics of reconstructed chromatin traces.

The pre-processing module organizes images by field of view (FOV) and by the type of probe imaged: DNA-FISH spots, nuclear/ oligopaint library masks, fiducial marks, or RNA expression. For each FOV, pyHiM first performs a projection and global registration using fiducial images acquired at each cycle as references (Fig. 1e). To improve the robustness of this step, we implemented a new method whereby the image is decomposed in blocks that are independently co-aligned. A polling step then determines the most popular global registration and applies it to the whole image (Fig. S2a). This step allows for a global correction of thermal drift and stage repeatability error even for cycles with fiducial images displaying local distortions. Samples such as embryos or tissues may often display local deformations during acquisition of different cycles which can not be taken into account by global registration algorithms. Thus, we developed a new local registration algorithm that optimizes 3D registrations locally to correct for 3D sample deformations (Figs. S2b, c).

The spot detection module performs segmentation and localization of DNA-FISH spots with sub-pixel accuracy of all sequential imaging rounds, using a combination of Deep Learning (DL)-powered spot segmentation followed by robust and automated 3D Gaussian fitting. 3D-DL segmentation is performed using a StarDist neural network ^18^ trained to robustly detect 3D-Point Spread Functions (PSF) in diverse sample types and illumination conditions. We obtained this network after extensive simulations of PSFs with different signal-to-noise ratios and inhomogeneous background levels. Next, based on the centroid position of each DL-mask, a robust 3D Gaussian fit of the intensity distribution is performed using Big-FISH (Figs. 1f and S3) ^19^.

The mask detection module segments nuclei in 3D using pre-trained StarDist neural networks models ^18^ (Fig. 1f). Other custom models based on StarDist or other popular architectures (*e*.*g*. Cellpose ^20^) can also be integrated via a plugin. Finally, DNA-FISH spots localized within the same mask are combined into chromatin traces, which are assigned a universally-unique identifier and tabulated in human-readable Enhanced Character-Separated Values ([ECSV](https://docs.astropy.org/en/stable/io/ascii/ecsv.html)) format (Figs. 1g and S4). Additional labels, based on RNA expression levels or spatial cell distribution, can be assigned to each single trace, allowing for cell/tissue-specific post-processing analysis (Fig. 1h).

Thanks to pyHiM’s modular architecture, each analysis step in the pipeline (registration, detection, tracing, *etc*) can be run independently. Users can tailor the analysis workflow according to their sample specificity, acquisition conditions and available computing resources (Fig. 2a). Intermediate results, such as unfiltered localizations or traces, are saved in ECSV format after each module execution, allowing the user to perform custom data validation or additional analysis. Finally, each module produces reports in human-readable markdown files with snapshot images illustrating the performance of the analysis for each cycle and FOV. This allows the user to efficiently assess the quality of the analyses and eventually fine-tune parameters to improve them (Figs. 2a-c and S2, S3). pyHiM can successfully analyze experimental data acquired from a variety of sample types, ranging from fly embryos to mouse and human tissues (Figs. 2d-e).

**Figure 2.**
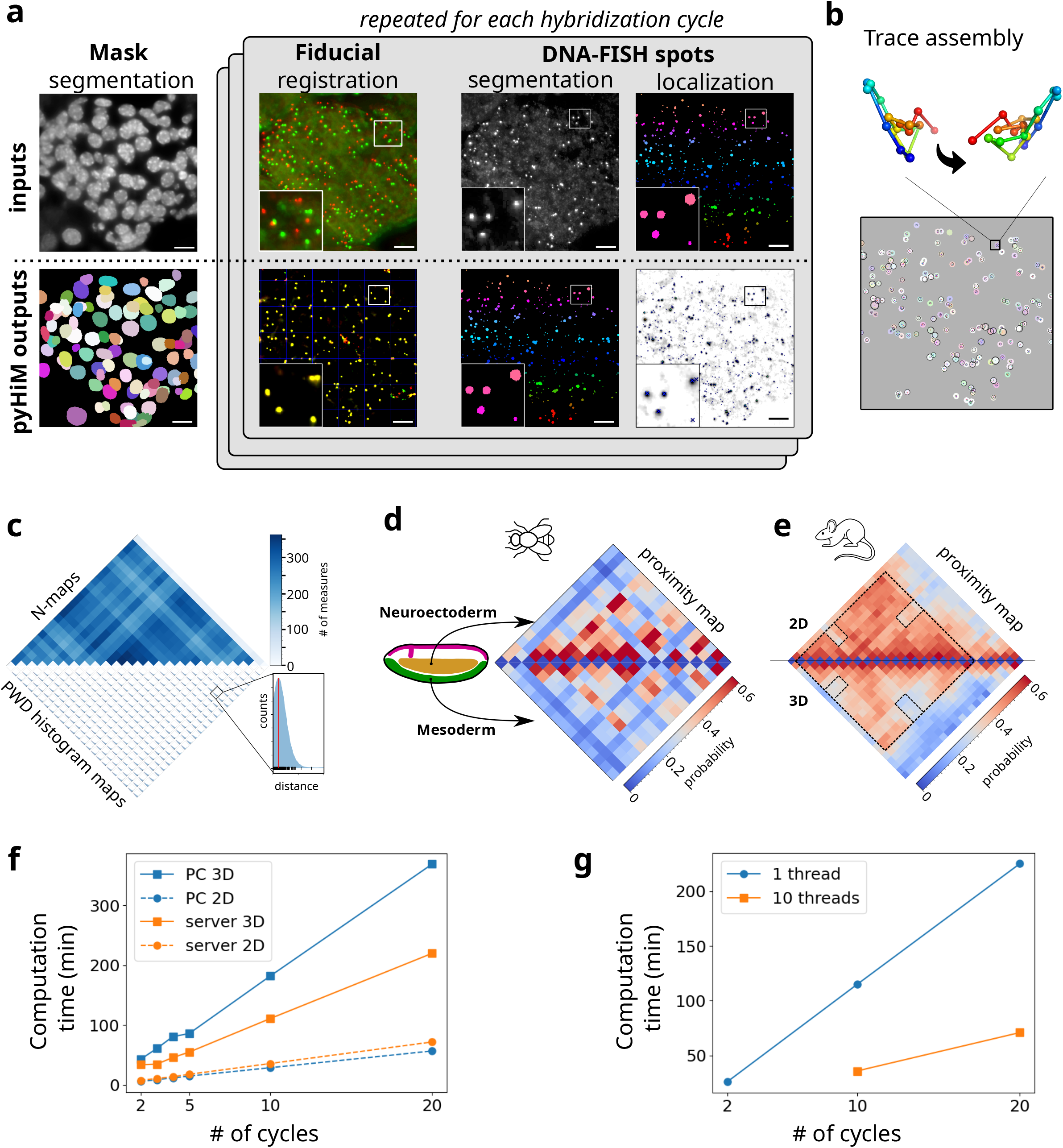
**a-** Illustration of a typical pyHiM analysis on mouse tissues: examples of raw data are shown in the top row and the most relevant pyHiM outputs are shown in the bottom row. From left to right, raw DAPI data are segmented to compute the 3D masks of each individual nucleus. Next, 2D & 3D registration of the fiducial is performed for each imaging cycle, and the quality of the correction can be quickly assessed based on the output image. Then, the localization of individual DNA-FISH spots is performed in two steps: first, a 3D mask of each DNA-FISH spot is computed using deep learning. Then, using the mask position as a reference, the sub-pixel localization of the spot is inferred using big-FISH. Scalebars = 8µm. **b-** Chromatin tracks are calculated by combining all individual DNA-FISH spot localizations detected within the same nuclear mask. Each individual trace represents a snapshot of the locus conformation within a single cell (see reconstruction with two different orientations). **c-** Data quality assessment: (top) The N-matrix represents the number of times that each pair of DNA loci was detected in the dataset, and is indicative of their detection efficiency. (Bottom) The distribution of pairwise distances between DNA-FISH spots in the same chromatin trace is plotted to ensure that there is no major error in the analysis (detection threshold, etc.). **d-** Traces computed by pyHiM were sorted based on RNA expression profiles in NC14 fly embryos and assigned to specific cell types (e.g. mesoderm vs. neuroectoderm). Specific long-range interactions and chromatin organization are observed for each cell type. **e-** Fast 2D analysis based only on the projected 3D data is used to optimize parameters and test data quality. An example from mouse tissue data shows the pairwise distance maps computed using 2D (top) and 3D (bottom) analysis. The 2D map captures most of the features that characterize the conformation of the studied locus. **f-** Comparison of pyHiM execution times for different number of cycles, and for a desktop computer (Intel(R) Core(TM) i7-8700 CPU @ 3.20GHz, CPUs: 12, Cores: 6, Threads per core: 2, Memory: 16Gb) or a multi-threaded server (AMD EPYC 7702 64-Core Processor 3.34GHz, CPUs:256, Cores: 128, Threads per core: 2, Memory: 512Gb). **g-** Performance of pyHiM using single-threaded or DASK-powered multi-threading.

pyHiM also offers a number of additional features that facilitate data formatting, result display, and post-processing. For instance, DNA-FISH spot detection efficiency and maps of the pairwise distance (PWD) distributions between DNA-FISH spots from different cycles (Fig. 2c), or proximity frequency matrices for specific cell-types (Fig. 2d). Another important feature of pyHiM is its ability to perform rapid analysis in 2D (Fig. 2e). In this mode, pyHiM projects signals from DNA-FISH spots and masks in 2D, and performs registration, segmentation, spot localization, and tracing in 2D. Contact maps computed using the 2D pipeline show all the relevant features of 3D maps (long-range contacts, TADs, *etc*), but require ∼5x less computation time (Figs. 2e-f) and can therefore be used to quickly assess the quality of the acquired dataset before full 3D analysis.

Finally, a critical aspect of multiplexed DNA-FISH imaging is the amount of data generated, typically ∼1-3 Tb per experiment depending on the number of cycles and the number of FOVs. To handle and analyze such large volumes of data in a reasonable time, we have implemented a parallelization mode based on the [Dask](https://www.dask.org/) Python package. For this, pyHiM analyzes data associated with different hybridization cycles in parallel, while keeping the technical aspects transparent to the user, leading to a drastic shortening in computation time (Figs. 2f-g). Conveniently, a reporting web-server based on [Bokeh](https://bokeh.org/) can be launched to monitor analysis status and performance in real-time (Fig. S4c). As a result, pyHiM can run indifferently on a laptop or an HPC cluster and be tuned according to the technical specificities of both (e.g. number of CPUs, available memory, availability of GPUs, *etc*).

In summary, we describe pyHiM, a modular, user-friendly, well-documented tool for chromatin tracing analysis based on sequential DNA-FISH imaging. pyHiM can be used to analyze data produced by Hi-M or by other spatial genomics methods. Thus, we envision that the adoption of pyHiM will enable the growth of a new user community for this active field of research. Indeed, as data acquisition and sample preparation become standard and even commercially available, a final bottleneck for widespread adoption will be the availability of flexible image analysis tool boxes dedicated to chromatin tracing. Thus, a well-tested and user-friendly analysis pipeline such as pyHiM will be key to break barriers to adoption of spatial genomics by users and microscopy facilities, to promote transparent image analysis pipelines in the field, and to create a large user community to accelerate discoveries and new developments. The modularity, open-source nature, and extensive developer documentation of pyHiM were purposefully designed to promote collaborative developments, to standardize and benchmark image analysis practices, and to facilitate reuse of existing algorithms to implement analysis tools for novel technologies in the blooming field of spatial genomics.

## Methods

### Inputs

The two minimal inputs of pyHiM are: a dictionary of parameters (*infoList*.*json*) and a list of images to process. *infoList*.*json* contains acquisition parameters (e.g. pixel size), file formatting parameters (e.g. regular expression to decode filenames), and all the parameters that are required for the execution of each module in pyHiM. For detailed information on the *infoList*.*json* parameter file, please refer to our online resource: [Input Parameters](https://pyhim.readthedocs.io/en/latest/reference/infoList_comprehension.html). Input images can be of two types: DNA-FISH spots for a given cycle, and masks used for tracing. The latter can be either nuclear masks (e.g. from DAPI labeling) or from a cycle where the whole oligopaint library is labeled and imaged at once. Both DNA-FISH and mask images must be accompanied by a corresponding fiducial image used for registration (see *Registration* section below). Images are assumed to be in the universal and non-proprietary [TIFF](https://docs.fileformat.com/image/tiff/) format. Use of deconvolved images is recommended but not compulsory.

### Projection

We developed a tool for image reprojection (module: *makeProjections*). This step is necessary for lateral global drift alignment (see *Registration* section below) and for the rapid visual inspection of input files. Sum and maximum projections are implemented and configured through the *infoList*.*json* parameters file. We recommend the former for masks and the latter for DNA-FISH images. makeProjections allows for the manual selection of the z-range, and implements an automatic algorithm to robustly retrieve the in-focus plane. Briefly, this method estimates the optimal in-focus plane by calculating the maximum of the laplacian of the intensity profile along the z-axis. The calculation is performed block-by-block to take into account local variability and sample drift. More details on the methods and the execution of this module can be found in the online description of the *[makeProjections](https://pyhim.readthedocs.io/en/latest/user_guide/modules/preprocessing/make_projections.html)* module.

### Registration

We implemented two registration methods to obtain automatic and robust global and local realignments. The *alignImages* module performs global realignments by registering the 2D z-reprojected fiducial images using 2D cross-correlation. This method, however, can be unreliable when fiducial images contain impurities that vary between cycles. To solve this, we developed a second algorithm (*alignByBlock*) that uses block-by-block decomposition to determine the best registration for each block. This calculation is followed by a polling operation that retrieves the most satisfactory global registration. This second method is highly robust to impurities. More details on the methods and the execution of this module can be found in our online description of the *[alignImages](https://pyhim.readthedocs.io/en/latest/user_guide/modules/preprocessing/align_images.html)* module. Once registrations for each cycle are processed, the module *[appliesRegistrations](https://pyhim.readthedocs.io/en/latest/user_guide/modules/preprocessing/applies_registrations.html)* re-interpolates 2D images of DNA-FISH spots and masks to provide a visual input of the performance of global registrations for each hybridization cycle.

Biological samples can display local deformations (typically in the hundreds of nm range) during the long-term acquisition times of a HiM dataset. These distortions cannot be properly corrected by global 2D realignment routines. To tackle this issue, we developed a new registration method that performs local 3D registration. In this method, images are first globally realigned in 2D. Next, fiducial images are decomposed in 3D blocks and each block is realigned by 3D cross-correlation and re-interpolation. The resulting local block corrections are stored as an ASTROPY table ^21^ that is used by the *register_localizations* module (see section below). More details on the methods and the execution of this module can be found in our online description of the *[alignImages3D](https://pyhim.readthedocs.io/en/latest/user_guide/modules/preprocessing/align_images_3d.html)* module.

### Segmentation and detection

Three different modules were built to deal with the segmentation and detection of DNA-FISH spots and masks. First, we developed a module for the segmentation and localization of masks and sources in 2D (module: *SegmentMasks*). Mask and DNA-FISH images are segmented using startdist with pre-trained networks. Segmented objects are filtered by size and shape, while merged objects are split using the watershed algorithm. DNA-FISH spots are fitted using the highly efficient DAOStarFinder algorithm from photutils ^23^, and post-processed using *filter_localizations*.

Second, we developed a module specifically designed to segment masks in 3D (module: *segmentMasks3D). segmentMasks3D* relies on deep-learning segmentation using a network that we trained specifically to robustly segment nuclei in 3D with stardist ^22^. Other DL segmentation tools, such as cellpose ^20^, can be used to further increase the flexibility of mask segmentation for different biological samples. *segmentMasks3D* then post-processes 3D masks by size and shape filtering, and applies a watershed algorithm to split merged masks. The output of *segmentMasks3D* is a 3D labeled image used by the *build_traces* module to group localizations into single chromatin traces (see *Tracing* section below). More details on the methods and the execution of this module can be found in our online description of the *[segmentMasks3D](https://pyhim.readthedocs.io/en/latest/user_guide/modules/identification/segment_masks_3d.html)* module.

Finally, we developed a module for the segmentation and localization of DNA-FISH spots (module: *segmentSources3D*). *segmentSources3D* segments DNA-FISH spots by using a stardist DL network trained to detect PSFs in 3D. This network was optimized by training the DL network on simulated data displaying large variations in signal-to-noise ratios, local background inhomogeneities, and intensity levels. After segmentation, *segmentSources3D* fits the intensity distributions within DNA-FISH spot masks with a 3D-Gaussian model using non-linear regression with functions from Big-FISH ^19^. The output *segmentSources3D* is an ASTROPY table containing the *xyz* coordinates, identities, and properties of all the localizations. Localizations with low intensities are filtered in post-processing using the module *filter_localizations*. A final step before tracing involves the application of local registrations to the localization tables obtained from *segmentMasks* or from *segmentSources3D using the register_localizations module*. The DL networks trained for pyHiM are available from our [pyHiM OSF repository](https://osf.io/ugpyh/)

### Tracing

The final step involves the grouping of DNA-FISH spots belonging to the same chromatin fiber (module: *build_traces*). This is accomplished in an iterative manner by grouping together localizations that belong to each segmented object in a mask image (either from nuclei or from labeling the entire oligopaint library). The output of *build_traces* is a trace table in ASTROPY format where each trace is stamped with a universal unique identifier to enable the automatic merging of multiple trace tables while avoiding misattributions. More details on the methods and the execution of this module can be found in our online description of the *build_traces* module.

We developed several tools for post-processing of trace tables. *Trace_selector* finds traces that match specific morphological or gene-expression patterns by matching trace localization with user-provided 3D masks. *Trace_combinator* merges traces from different FOVs or different experiments. *Trace_filter* is a general tool for filtering traces that can remove specific barcodes from a trace table, remove duplicated localizations from single traces, and perform spatial filtering. *[Trace_analyzer](https://pyhim.readthedocs.io/en/latest/user_guide/modules/building_traces/trace_analyser.html)* analyzes a trace table to calculate the distribution in the number of barcodes detected per trace, the number of times each barcode appears in single traces, and the spatial clustering of traces.

Finally, we developed an algorithm that builds maps from trace tables (module: *build_matrix*). This tool produces conventional pair-wise median distance maps, relies on kernel-density estimators to accurately calculate the maximum of each distance distribution, and calculates proximity distance maps for user-specified threshold distances. *Build_matrix* produces N-maps which contain the number of localizations detected for each combination of barcodes, a diagnostic tool that is fundamental to determine the performance of an experiment and the robustness of detection for each barcode pair. More details on the methods and execution of this module can be found in our online description of the *[*build_matrix*](https://pyhim.readthedocs.io/en/latest/user_guide/modules/building_traces/build_matrices.html)* module.

### Code availability

The latest stable and development versions of pyHiM are publicly available at our Github repository: https://github.com/marcnol/pyHiM. The online documentation is available at: https://pyhim.readthedocs.io/en/latest/.

## Acknowledgments

This project was funded by the European Union’s Horizon 2020 Research and Innovation Program (EpiScope, Grant ID 724429, M.N.). We acknowledge the Bettencourt-Schueller Foundation for their prize ‘Coup d’élan pour la recherche Française’, and the France-BioImaging infrastructure supported by the French National Research Agency (grant ID ANR-10-INBS-04, ‘‘Investments for the Future’’). Part of this work was funded by the cloudFISH project from France-BioImaging.

---

**Supplementary Figure 1**

**a-** List of all the python Packages used by pyHiM and their versions.

**b-** Graphical user interface for pyHiM parameter setting routine.

**Supplementary Figure 2**

**a-** pyHiM first performs a 2D registration on projected 3D images: Example of two superimposed fiducial images before (left) and after (middle) registration. The registration can be performed globally or locally. In the latter case, the image is divided into an array of 8x8 blocks and the registration is computed separately for each block. The block correction map (right) shows the shift along the x-axis applied to the registered image (in pixels, 1 pixel is 105 nm). Scalebars = 25 µm.

**b-** The 2D global registration can be affected by local sample deformation, and a 3D block-by-block routine is used to further improve the correction. Again, a local correction is computed by dividing the 3D images into an array of 16 x 16 x 16 blocks. A quick assessment of the correction quality can be made based on the overlaid images. Scalebars = 25 µm.

**c-** Average block correction maps for 3D registration are used to check for errors or unexpected deformation of the sample during the experiment.

**Supplementary Figure 3**

**a-** DAPI-stained nuclei are imaged in a Drosophila embryo (left) and segmented using StarDist (right). Scalebar = 25µm.

**b-** Using a library of oligonucleotides targeting a large genomic region, bright 3D fiducial images are acquired in mouse tissue (left). Using custom trained StarDist models, the images are segmented and individual masks are computed for each fiducial. Scalebar = 25µm.

**c-** For each round of acquisition, individual DNA-FISH images are analyzed and sub-pixel localizations are computed using big-FISH. Each detection is assigned a unique ID and saved together with its x, y, z localizations and intensity in a. *dat* file. The table summarizes all the parameters available for each detection.

**d-** The distribution of peak intensity is plotted against spot localizations along the z-axis. Such data can be used to further optimize the intensity threshold used for detection and check for illumination artifacts.

**e-** Example of detection output. Each individual detections are represented by a spot of color overlay with the maximum intensity projection of the original image (in inverted grayscale). Scalebars = 25µm.

**Supplementary Figure 4**

**a-** 3D representation of example chromatin traces.

**b-** Format of the chromatin table output file.

**c-** Snapshot of Bokeh server displaying the advancement of a pyHiM execution in parallel mode.

