## Supplementary Figures for "pyHiM, a new open-source, multi-platform software package for spatial genomics based on multiplexed DNA-FISH imaging"

### Supplementary Figure 1

**a**

**pyHiM dependencies**

| Library | Version |
|---|---|
| big-fish | >= 0.6.2 |
| astropy | >= 4.3.1 |
| csbdeep | >= 0.6.3 |
| opencv-python | >= 4.5.3.56 |
| dask[distributed] | >= 2021.10.0 |
| matplotlib | >= 3.5.0 |
| numba | >= 0.54.1 |
| numpy | >= 1.19.5 |
| photutils | == 1.1.0 |
| pylab | >= 1.3.2 |
| pympler | >= 1.0.1 |
| roipoly | >= 0.5.3 |
| scipy | >= 1.7.3 |
| scikit-image | >= 0.18.3 |
| scikit-learn | >= 1.0.2 |
| stardist | >= 0.7.3 |
| tifffile | >= 2021.7.2 |
| tqdm | >= 4.62.3 |

**b**

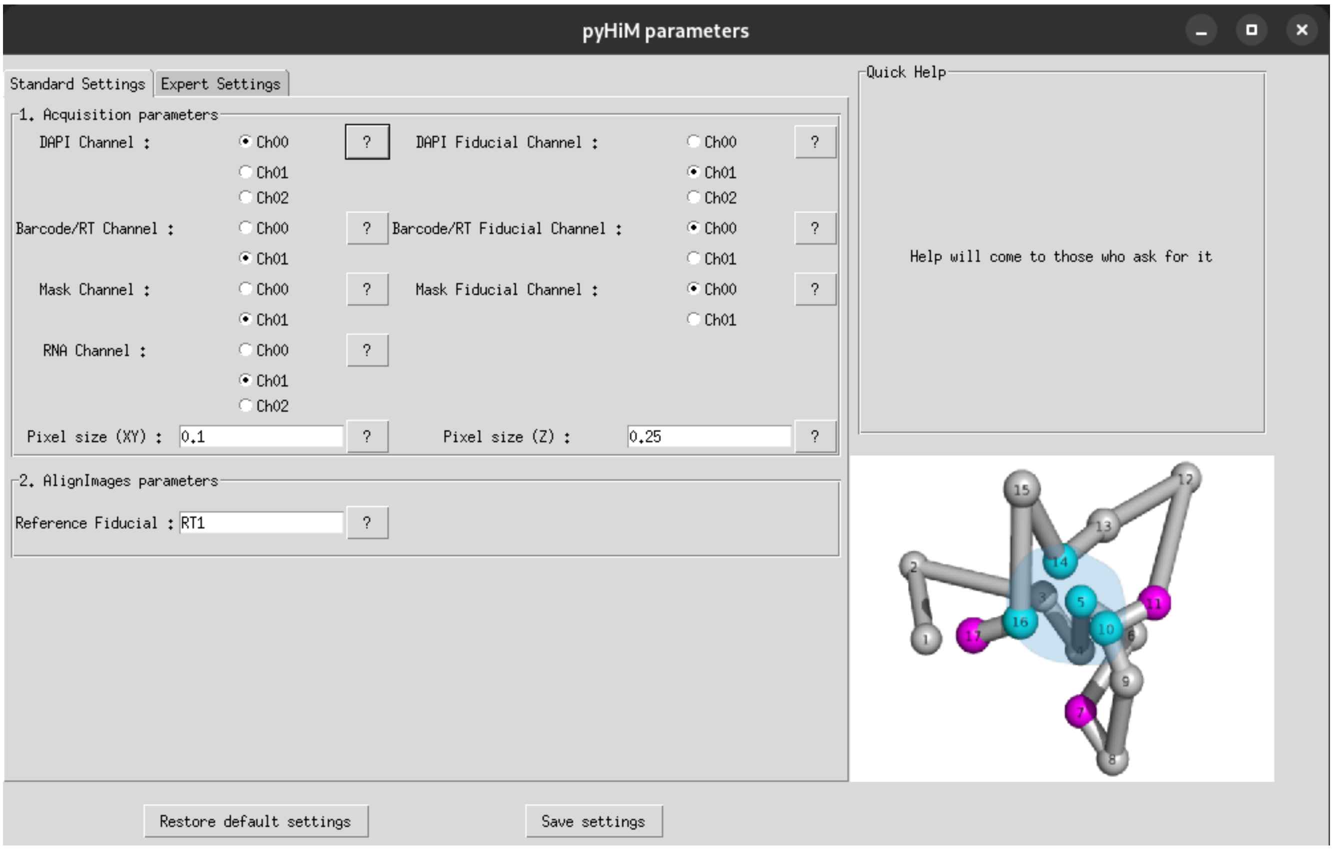

### Supplementary Figure 2

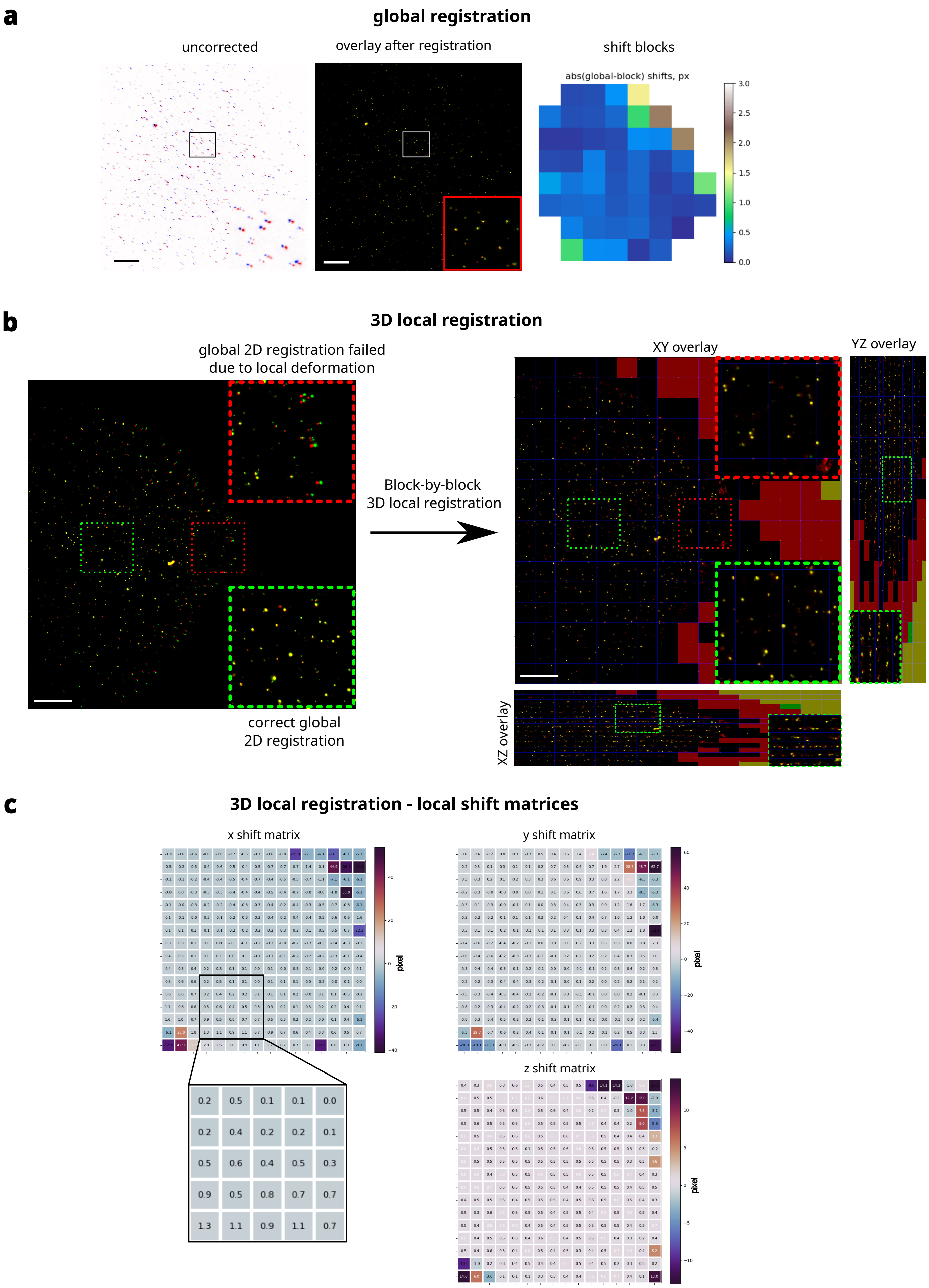

### Supplementary Figure 3

**a** **segmentation DAPI masks**

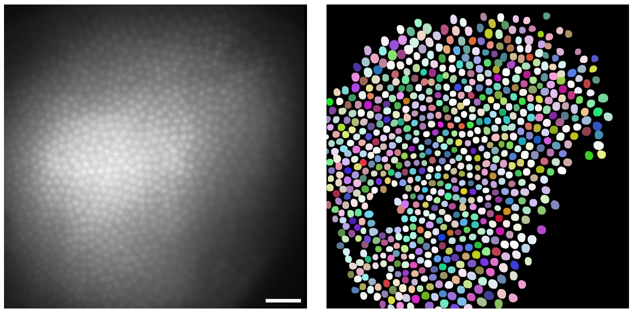

**b** **segmentation entire library mask**

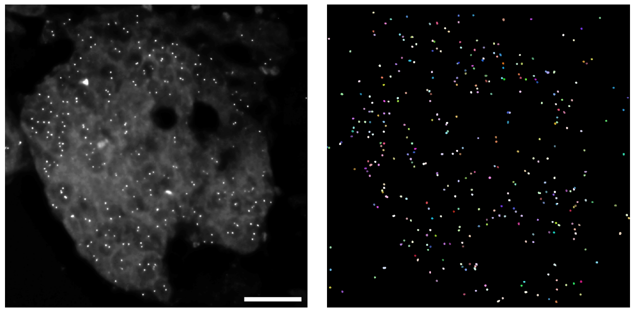

**c** **localizations table**

| Column | Description |
| --- | --- |
| Buid | Unique identifier of spot |
| ROI # | Region of interest number |
| CellID # | Cell identifier number |
| Barcode # | DNA-FISH probe number |
| id | Unique spot identifier per barcode |
| zcentroid | Localization along z-axis (in plane) |
| xcentroid | Localization along x-axis (in px) |
| ycentroid | Localization along y-axis (in px) |
| sharpness | Ratio between max intensity and mean background |
| roundness1 | Object roundness based on symmetry |
| roundness2 | Object roundness based on Gaussian fits |
| npix | Number of pixels in the Gaussian kernel array |
| sky | Input background intensity (default is 0) |
| peak | Maximum pixel value of the spot |
| flux | Peak density divided by the detection threshold |
| mag | -2.5 * log10(flux) |

**d** **localization statistics**

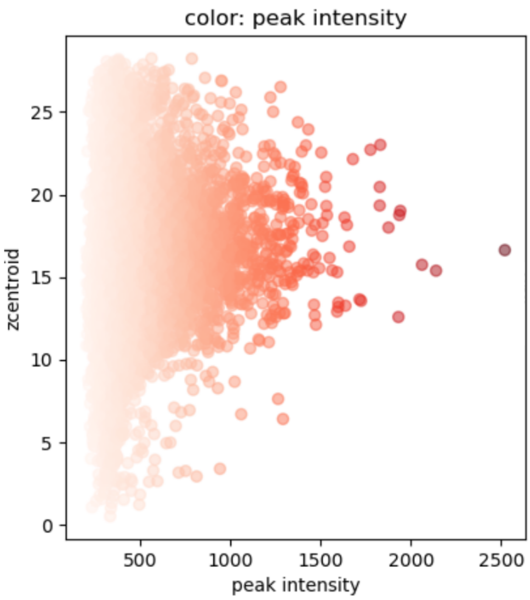


**e** **barcode masks and localizations**

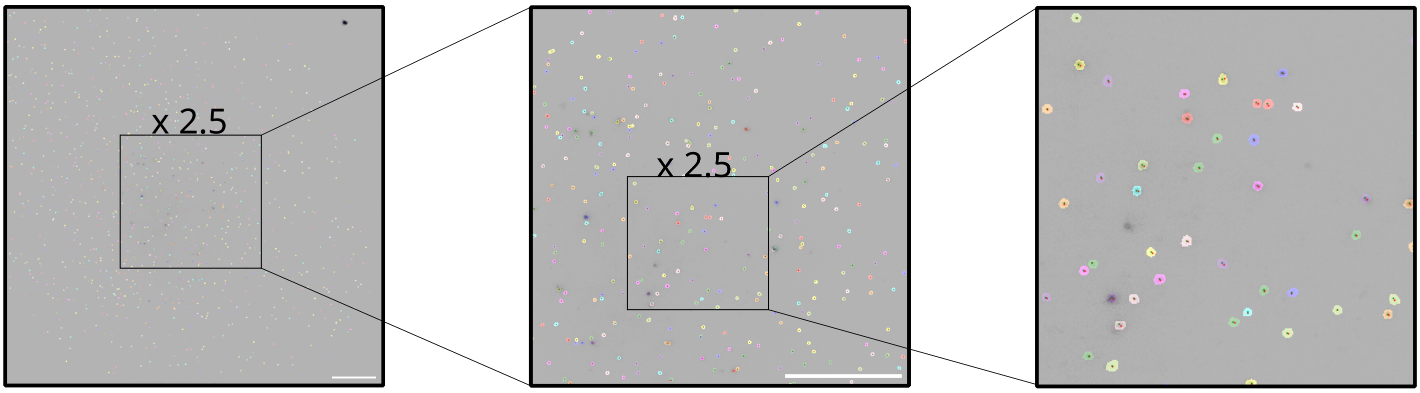

### Supplementary Figure 4

**a** example traces

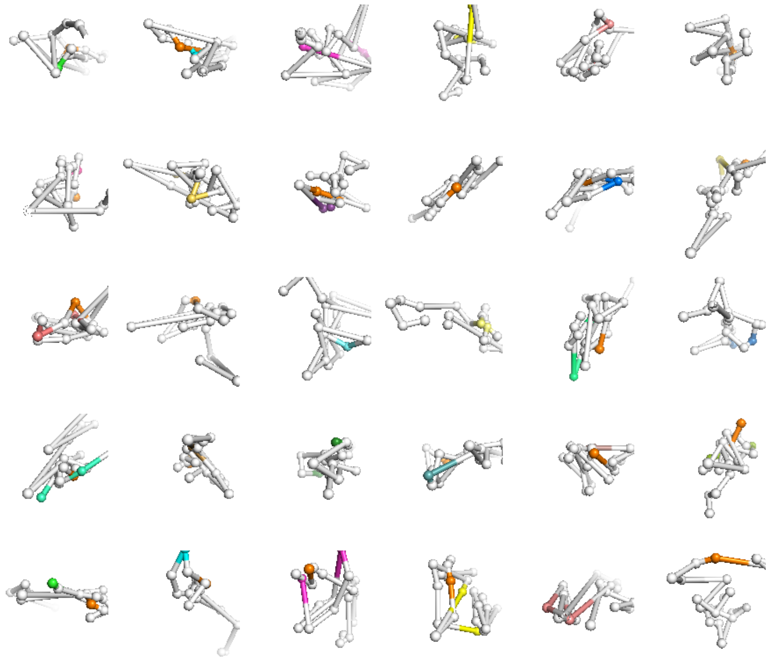

**b** trace table

| Column | Description |
|---|---|
| Spot_ID | Spot unique identifier |
| Trace_ID | Trace unique identifier |
| x | x coordinate (in nm) |
| y | y coordinate (in nm) |
| z | z coordinate (in nm) |
| Chrom | Chromosome |
| Chrom_Start | Sequence start (in bp) |
| Chrom_End | Sequence end (in bp) |
| ROI # | Region of interest number |
| Mask_id | Mask identifier |
| Barcode # | DNA-FISH probe number |
| label | Optional label to classify trace |

**c** online Bokeh server

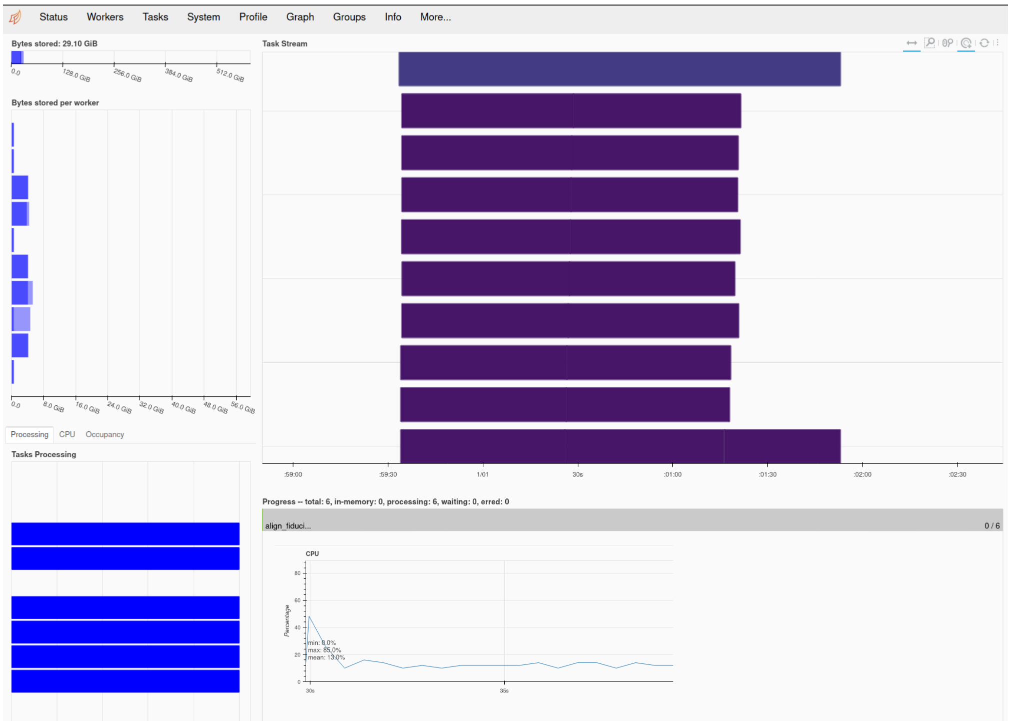


| name | address | nthreads | cpu | memory | limit | memory % | managed | unmanage | unmanage | spilled | # fds | read | write |
|---|---|---|---|---|---|---|---|---|---|---|---|---|---|
| Total (10) | | 10 | 980 % | 9.2 GiB | 596.0 GiB | 1.5 % | 0.0 | 7.6 GiB | 1.6 GiB | 0.0 | 249 | 442 KiB | 2 MiB |
| 0 | tcp://127.0 | 1 | 102 % | 851.8 MiB | 59.6 GiB | 1.4 % | 0.0 | 720.8 MiB | 131.0 MiB | 0.0 | 29 | 19 KiB | 19 KiB |
| 1 | tcp://127.0 | 1 | 102 % | 1.5 GiB | 59.6 GiB | 2.5 % | 0.0 | 774.9 MiB | 744.7 MiB | 0.0 | 24 | 30 KiB | 30 KiB |
| 2 | tcp://127.0 | 1 | 96 % | 832.0 MiB | 59.6 GiB | 1.4 % | 0.0 | 767.1 MiB | 64.9 MiB | 0.0 | 24 | 19 KiB | 19 KiB |
| 3 | tcp://127.0 | 1 | 104 % | 944.1 MiB | 59.6 GiB | 1.5 % | 0.0 | 758.6 MiB | 185.5 MiB | 0.0 | 24 | 54 KiB | 367 KiB |
| 4 | tcp://127.0 | 1 | 3262 % | 868.0 MiB | 59.6 GiB | 1.4 % | 0.0 | 795.7 MiB | 72.3 MiB | 0.0 | 24 | 60 KiB | 373 KiB |
| 5 | tcp://127.0 | 1 | 2 % | 704.6 MiB | 59.6 GiB | 1.2 % | 0.0 | 703.7 MiB | 832.0 KiB | 0.0 | 28 | 60 KiB | 374 KiB |
| 6 | tcp://127.0 | 1 | 100 % | 835.2 MiB | 59.6 GiB | 1.4 % | 0.0 | 768.8 MiB | 66.4 MiB | 0.0 | 24 | 60 KiB | 373 KiB |
| 7 | tcp://127.0 | 1 | 4221 % | 800.1 MiB | 59.6 GiB | 1.3 % | 0.0 | 751.1 MiB | 48.9 MiB | 0.0 | 24 | 60 KiB | 373 KiB |
| 8 | tcp://127.0 | 1 | 109 % | 1.1 GiB | 59.6 GiB | 1.9 % | 0.0 | 783.9 MiB | 365.3 MiB | 0.0 | 24 | 19 KiB | 19 KiB |
| 9 | tcp://127.0 | 1 | 1703 % | 921.7 MiB | 59.6 GiB | 1.5 % | 0.0 | 920.9 MiB | 808.0 KiB | 0.0 | 24 | 61 KiB | 375 KiB |
